# The role of lipoteichoic acid in *Staphylococcus aureus* cell wall integrity

**DOI:** 10.1101/2025.01.16.633316

**Authors:** Amol Kanampalliwar, Majid Shah, Youngseon Park, Bohyun Jeong, Paul A. Lawson, Madeline Bell, Elizabeth Margaret Fesko, Anthony Sainato, Dominic Sainato, Suzanne Walker, Taeok Bae

## Abstract

In *Staphylococcus aureus*, lipoteichoic acid (LTA) is crucial for growth, cell division, osmoprotection, and beta-lactam resistance, yet its molecular mechanisms remain unclear. This study reveals that LTA binds to multiple proteins involved in cell wall processes and preserves cell wall integrity by regulating one of the LTA-binding proteins, ScaH. ScaH is a peptidoglycan hydrolase predicted to have N-acetylglucosaminidase and amidase/peptidase activities. LTA inhibits ScaH’s enzymatic activities by direct binding and represses *scaH* transcription by an unknown mechanism. During early growth, LTA is highly expressed and sequesters ScaH at the cell membrane, preventing ScaH activity in the cell wall. However, LTA expression decreases during the late growth phase, leading to ScaH translocation into the cell wall. This reduction in LTA coincides with increased wall teichoic acid (WTA) expression and cleavage of the LTA synthase LtaS. In the LTA-null mutant, ScaH inactivation restored peptidoglycan crosslinking, osmoresistance, and beta-lactam resistance both *in vitro* and *in vivo*. These findings suggest that LTA protects the cell wall by suppressing ScaH expression and activity while sequestering it during active growth. Additionally, the reciprocal expression patterns of LTA and WTA indicate an interconnected regulation of teichoic acids in *S. aureus*, with their roles likely depending on the growth phase.

**Importance:** *Staphylococcus aureus* is a significant human pathogen, with LTA playing a crucial role in cell viability, division, and maintaining cell wall integrity. Targeting LTA synthesis holds promise for the development of new therapeutics against *S. aureus*. Disruption of LTA synthesis leads to phenotypes indicative of compromised cell wall integrity, such as reduced crosslinking, osmosensitivity, and heightened beta-lactam sensitivity, although the precise underlying mechanisms remain unclear. Our research demonstrates that LTA contributes to cell wall integrity by regulating the expression, activity, and translocation of the peptidoglycan hydrolase ScaH. Additionally, LTA’s binding to the cell wall synthesis enzymes suggests a role in modulating cell wall integrity through these interactions. These findings significantly advance our understanding of LTA’s physiological functions and may serve as a foundation for the development of novel therapeutics targeting *S. aureus* and other Gram-positive pathogens.

## Introduction

Teichoic acids (TAs) are anionic polymer molecules found on the surface of Gram-positive bacteria. Two types of TAs exist: lipoteichoic acid (LTA) on the cell membrane and wall teichoic acid (WTA) on the cell wall (1, 2). The synthesis pathways and the chemical structures of TAs vary among bacteria (3). TAs play important roles in bacterial growth, morphology, virulence, and resistance to stresses and antimicrobials (1, 2), yet the molecular mechanisms underlying these functions remain largely unclear.

*Staphylococcus aureus* is a Gram-positive pathogen responsible for a wide range of diseases. The rise of multidrug-resistant strains, such as methicillin-resistant *S. aureus* (MRSA), has made treating staphylococcal infections increasingly difficult, driving the search for new therapeutic targets. TA synthesis pathways are considered promising targets (4), with LTA synthesis being attractive due to its essential role in staphylococcal growth (5–7).

In *S. aureus*, LTA is a polymer of phosphoglycerol, and its last synthesis steps involve three proteins: UgtP (also known as YpfP), LtaA, and LtaS (8, 9). UgtP produces the membrane anchor molecule diglucosyl diacylglycerol (Glc_2_-DAG) (10), which is flipped to the outer layer of the membrane by LtaA (9). LtaS then transfers phosphoglycerol from phosphatidylglycerol to the anchor molecule to complete LTA synthesis (8, 11). In laboratory growth conditions, the deletion of *ltaS* is lethal in most staphylococcal strains, and the survival of the *ltaS* mutant requires a low temperature (e.g., 30°C) or osmoprotectants (e.g., 7.5% NaCl and 40% sucrose) (12, 13) or compensatory mutations in *clpX*, *gdpP, sgtB,* and *mazE* (13–15). The *ltaS* mutant typically shows reduced peptidoglycan (PG) crosslinking, enlarged cell size, aberrant septum formation, osmosensitivity, and increased susceptibility to various antimicrobial agents, such as lysostaphin, vancomycin, and oxacillin (5, 12, 13, 15). These phenotypes suggest defective cell wall integrity, but the precise mechanisms underlying these defects remain unknown.

The cell wall is the critical structure in most bacteria, maintaining cell shape and withstanding intracellular turgor pressure. Its integrity is largely controlled by two opposing groups of enzymes: penicillin-binding proteins (PBPs)/SEDS (shape, elongation, division, sporulation) glycosyltransferases, which synthesize cell wall (16), and peptidoglycan hydrolases (PGHs), which degrade and remodel it (17). All *S. aureus* produce four PBPs (PBP1 - 4), with PBP2 being the only one containing both transpeptidase and glycosyltransferase domains. PBP2a is a β-lactam resistant transpeptidase encoded by a mobile genetic element and unique to MRSA (18, 19). For cell wall degradation and remodeling, *S. aureus* produces 18 PGHs, of which Atl is known to bind LTA (20, 21).

In this study, we investigated the role of LTA in cell wall integrity using a *ltaS* deletion mutant carrying a compensatory mutation. We found that LTA directly binds to two PBPs (i.e., PBP2 and PBP4) and three PGHs (i.e., Atl, ScaH, and SagA). Further analysis suggests that LTA binding to ScaH is central to the protective role of LTA in cell wall integrity.

## Results

### The essentiality of LTA could be mitigated by mutations in *cozEb*

The *ltaS* gene has been reported as nonessential in the MW2 strain, although a nonsynonymous SNP in *clpX* was suspected of contributing to the *ltaS* nonessentiality of the strain (5). We deleted *ltaS* in MW2 using the allele-replacement plasmid pCasSA-v2, an improved version of pCasSA (22) (Fig. S1). The deletion mutant was isolated at 30°C without osmoprotectants, and its successful deletion was confirmed by PCR and Western blotting (Fig. S2). As anticipated, MW2Δ*ltaS* exhibited near-normal growth at 37°C without osmoprotectants (Fig. 1A).

**Figure 1.**
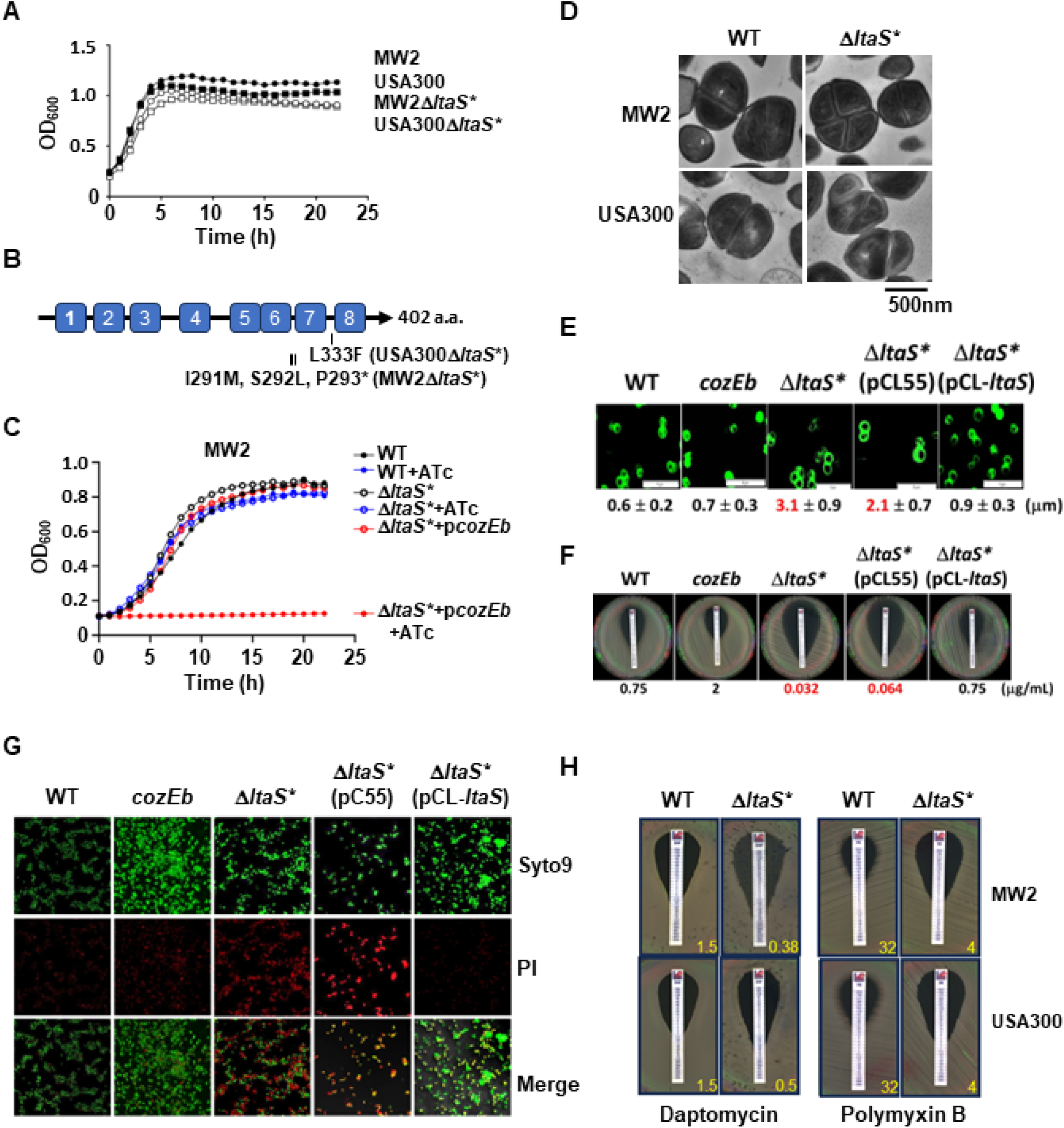
Characterization of Δ*ltaS*\*. **(A)** Growth of Δ*ltaS*\* in TSB at 37°C. **(B)** CozEb suppressor mutations found in Δ*ltaS\**. Numbers indicate transmembrane domains, and each vertical bar represents a mutation site. MW2Δ*ltaS\** has three mutations (T>TG, C>A, C>AG) between A290 and P293, resulting in I291M, S292L and P293* mutations. **(C)** Effect of *cozEb* complementation on the growth of MW2Δ*ltaS*.* Test strains were grown in TSB at 37°C with or without anhydrotetracycline (ATc, 100 ng/mL). **(D)** Cell division defects of Δ*ltaS\**. Cells were grown in TSB until the exponential growth phase and observed by TEM. **(E)** Effect of Δ*ltaS\** mutation on the cell size of USA300 strain, assessed by confocal microscopy. Cells at the exponential growth phase were stained with FITC-labeled vancomycin and observed by confocal microscopy. The quantification graph is in Fig. S5. **(F)** Oxacillin susceptibility of USA300Δ*ltaS\**. MIC values are shown under each image. **(G)** Osmosensitivity of USA300Δ*ltaS\**. Cells were grown to the exponential growth phase, collected, suspended in water, and incubated for 6 h. PI, propidium iodide. The quantification graph is in Fig. S6B. **(H)** Sensitivity of USA300Δ*ltaS\** to cell-membrane-targeting antibiotics. MIC values are shown at the bottom right of each image.

Unexpectedly, when we deleted *ltaS* in USA300 using the same method (Fig. S2), the resulting USA300Δ*ltaS* also displayed almost normal growth at 37°C (Fig. 1A), contrary to the previous report that *ltaS* is essential in this lineage (5). Whole-genome sequencing of both MW2Δ*ltaS* and USA300Δ*ltaS* revealed compensatory mutations in *cozEb*, encoding a cell division protein shown to interfere with LTA biosynthesis (Fig. 1B and S3) (23, 24). When *cozEb* was expressed from an anhydrotetracycline (ATc)-inducible promoter in a multi-copy plasmid, neither mutant was able to grow at 37°C (Fig. 1C and S4), indicating that the *ltaS* gene is essential even in MW2 and that the *cozEb* mutations compensate the lack of LTA under physiological conditions. Due to the presence of compensatory mutations, we named these mutants as Δ*ltaS*\*.

### Δ*ltaS\** mutants maintain the phenotypes expected from Δ*ltaS*

Despite the compensatory mutations, the Δ*ltaS\** mutants exhibited phenotypes consistent with those expected from *ltaS* deletion mutants, such as defects in cell division (Fig. 1D), increased cell size (Fig. 1E and S5), decreased resistance to oxacillin (Fig. 1F), increased osmosensitivity (Fig. 1G and S6), and increased thermosensitivity (Fig. S7) (12, 13). Notably, the *cozEb* mutation alone did not induce those phenotypes (*cozEb* in Fig. 1E-G). Furthermore, introducing a single copy of *ltaS* restored normal phenotypes (Fig. 1E-G and Fig. S5-S7), demonstrating that those phenotypes are due to the *ltaS* deletion, not the compensatory mutation.

Given that the *ltaS* deletion mutant is not viable under physiological conditions, we used the Δ*ltaS\** strains to investigate further the roles of LTA in cell wall integrity.

### The *ΔltaS\** mutants are more susceptible to membrane-attacking antibiotics

The *ugtP* and *ltaA* mutants are more susceptible to daptomycin, which targets cell membranes (5). Interestingly, the *ΔltaS\** strains also demonstrated heightened sensitivity to daptomycin, as well as to polymyxin B, another cell-membrane targeting antibiotic (Fig. 1H). These results led us to investigate whether the fatty acid composition of the cell membrane was altered in the *ΔltaS\** strains and found no significant changes (Fig. S8), indicating that the increased sensitivity to those membrane-targeting antibiotics is likely due to alterations in the cell wall, not in the membrane.

### LTA binds PBPs and PGHs

Next, we identified LTA-binding proteins by affinity chromatography. LTA was crosslinked to an epoxide resin and the membrane fraction of USA300 WT was applied and eluted with high salt. Since LTA is modified with D-alanine, we generated an alanine affinity column for a control. From the LTA affinity column, several proteins between 50 - 80 kDa were eluted explicitly (Fig. 2A). Mass spectrometry analysis of those protein bands identified three membrane proteins (PBP2, FtsH, and DivIB) and two secreted PGHs (Atl and ScaH) (Fig. 2B). Atl is known to bind LTA (21), validating our approach. ELISA assays with recombinant candidate proteins and other PBPs & PGHs showed that LTA binds to two PBPs (PBP2, PBP4) and three PGHs (Atl, ScaH, and SagA) (Fig. 2C). LTA also showed minimal binding to PBP2a and SceD (Fig. 2C). These results indicate that LTA might protect cell wall integrity by interacting with PBPs and PGHs.

**Figure 2.**
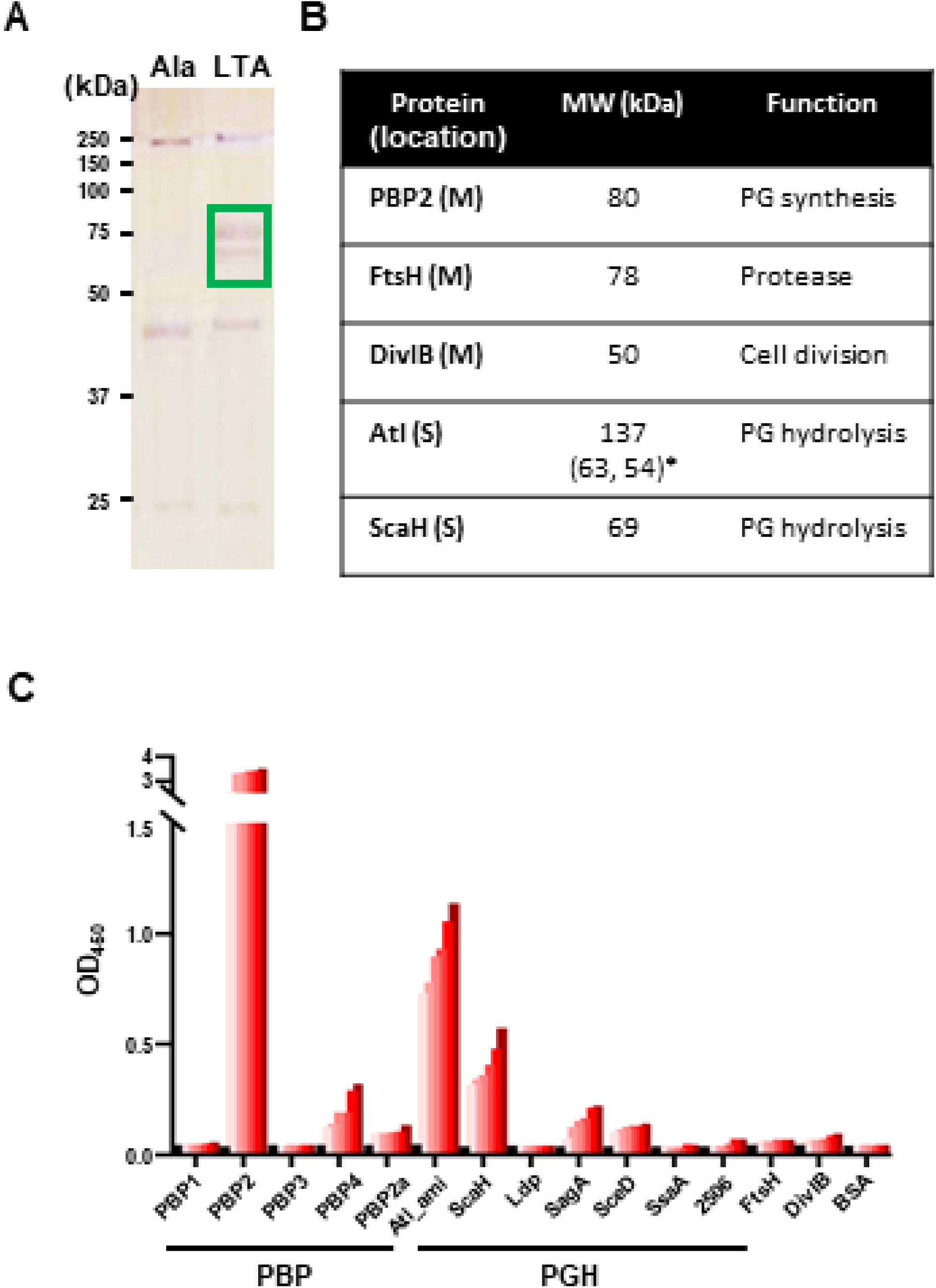
Identification of LTA binding proteins. **(A)** Proteins eluted from the alanine-affinity column (Ala) and the LTA-affinity column (LTA). The green box highlights the protein bands sent for mass spectrometry (MS) analysis. **(B)** Five candidate proteins identified by MS analysis. The asterisk (*) indicates the molecular weights of the processed Atl fragments. PG, peptidoglycan; M, membrane; S, secreted. **(C)** Confirmation of LTA binding by the candidate proteins using ELISA. Other penicillin-binding proteins (PBPs) and peptidoglycan hydrolases (PGHs) were also included to assess binding specificity. Proteins were immobilized on the ELISA plate, various amounts of LTA were added, and bound LTA was measured with an anti-LTA antibody. Atl_ami, Atl amidase.

### LTA synthesis suppresses *scaH* transcription

Intriguingly, *scaH* transcription was highly increased during the exponential growth phase in *ΔltaS\** strains (Fig. 3A and S9). In the stationary growth phase, however, upregulation of *scaH* was not observed (Fig. 3A and S9). The promoter-*lacZ* fusion assay recapitulated the qRT-PCR results (Fig. 3B). The *scaH* promoter activity was highly upregulated in *ΔltaS\** only in the exponential growth phase; in the stationary growth phase, *ltaS*-deletion did not affect *scaH* promoter activity. In the WT strain, *scaH* promoter activity was increased in the stationary growth phase, compared with the exponential growth phase (WT in Fig. 3B). Western blotting further confirmed the transcriptional analyses (Fig. 3C). These findings demonstrate that *scaH* promoter activity is inhibited by LTA synthesis during active growth, and it is also regulated in a growth-dependent manner in WT.

**Figure 3.**
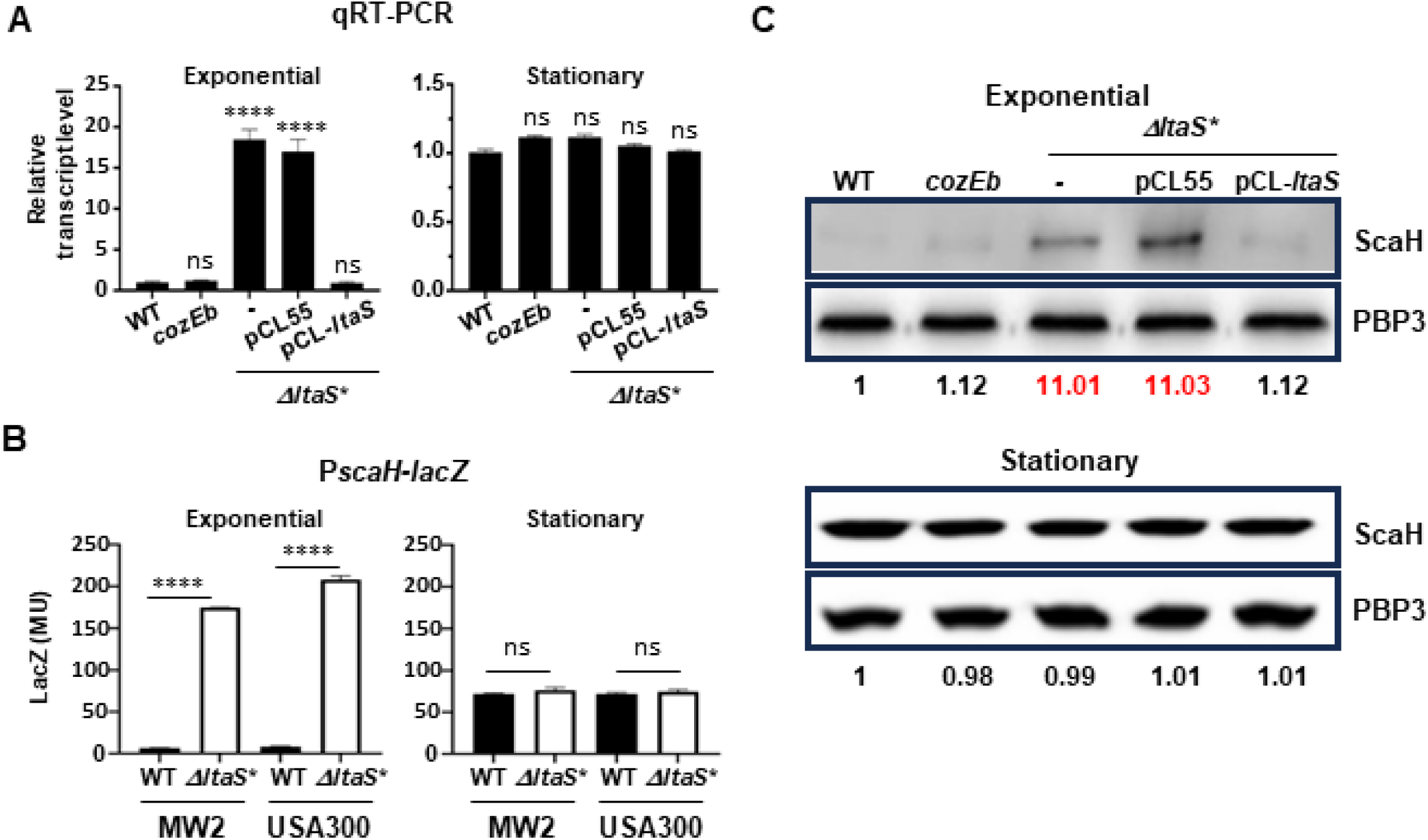
Suppression of *scaH* expression by LTA synthesis. **(A)** qRT-PCR analysis of *scaH* transcript levels. WT, wild type; coz*Eb*, the *cozEb* mutant; -, no plasmid; pCL55, vector control; pCL-*ltaS,* complementation plasmid. **(B)** Promoter-*lacZ* assay for *scaH* promoter activity. The *lacZ* gene was fused to the *scaH* promoter (P*scaH*), and its expression was measured during the exponential and stationary growth phases. MU, Miller unit. **(C)** Western blot analysis of ScaH expression. PBP3 was used as a loading control. Relative ScaH:PBP3 ratios are shown under each lane. ns, not significant; ****, *p* < 0.0001 by two-tailed unpaired t-test.

### LTA inhibits ScaH activity

ScaH has two possible enzymatic domains: an N-acetylglucosaminidase (NAGase) domain and an amidase/peptidase (CHAP) domain (Fig. 4A). We determined the LTA-binding regions of ScaH by ELISA with ScaH fragments and found that LTA binds to the central region of ScaH containing the NAGase domain (ΔN222ΔCHAP in Fig. 4A and 4B). To examine whether LTA binding affects cell wall degradation by ScaH, we carried out the Remazol brilliant blue (RBB)-release assay (25). RBB-labeled cell wall was incubated with ScaH alone or a ScaH/LTA mix, and the dye released by the cell wall degradation was measured by monitoring optical density (OD_595)_. As shown, LTA suppressed dye release in a time-dependent manner (Fig. 4C). These results demonstrate that LTA binding inhibits the cell wall degradation activity of ScaH.

**Figure 4.**
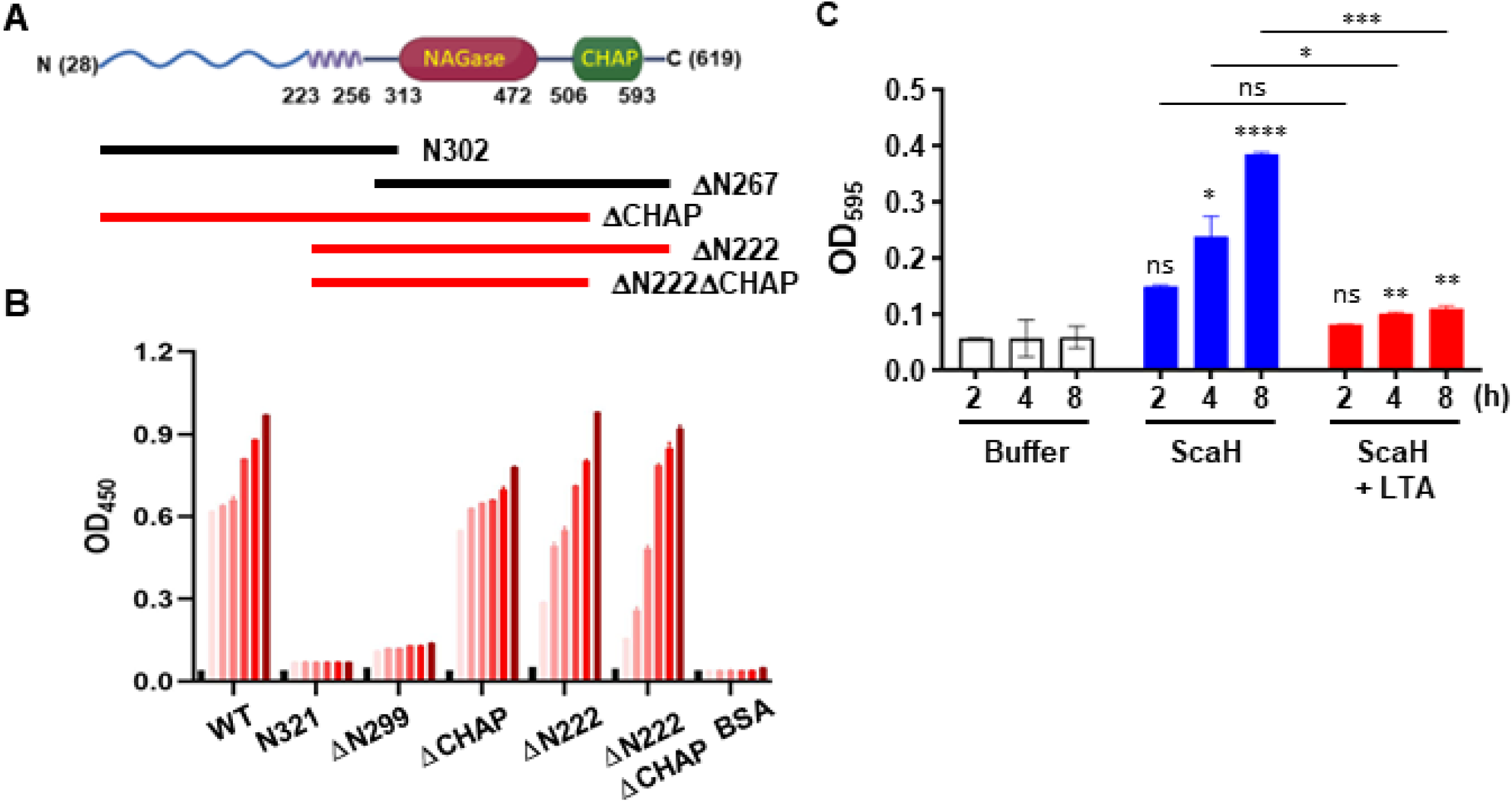
Inhibition of ScaH activity by LTA binding. **(A)** Domain structure of ScaH. The ScaH fragments are indicated with thick lines. Black indicates no binding, while red indicates binding. Numbers represent amino acid positions. NAGase, N-acetylglucosaminidase domain; CHAP, cysteine, histidine-dependent amidohydrolases/peptidases domain. **(B)** LTA binding of ScaH fragments assessed by ELISA. Proteins were immobilized on the ELISA plate, various amounts of LTA were added, and bound LTA was measured with an anti-LTA antibody. **(C)** Remazol brilliant blue (RBB)-release assay. Purified peptidoglycans were coated with RBB and treated with ScaH or a ScaH + LTA mix. Released dyes were measured by monitoring optical density at 595 nm every 2 h. ns, not significant; *, *p* < 0.05; **, *p* < 0.01; ***, *p* < 0.0005; ****, *p* < 0.0001 by two-tailed unpaired t-test.

### LTA and WTA are produced reciprocally in a growth-dependent manner

Since the *scaH* promoter activity was suppressed by LTA synthesis but activated in the stationary phase (Fig. 3B), we measured LTA expression throughout bacterial growth. LTA was highly expressed during the early growth phase and subsequently downregulated as the bacteria entered the late growth phase (Fig. 5A and 5B). This downregulation coincided with the depletion of intact LtaS (Fig. 5B). Intriguingly, WTA displayed a reciprocal expression pattern: Its expression was minimal during the early growth phase but upregulated during the late growth phase (Fig. 5B). Moreover, the translocation of ScaH followed the LTA expression. In the WT strain, ScaH was localized in the membrane during the early growth phase; however, during the later growth phase, ScaH expression increased, and ScaH was found in the cell wall (Fig. 5C). Conversely, in the Δ*ltaS*\* mutant strain, ScaH was constitutively produced at a higher level than WT and consistently located in the cell wall regardless of the bacterial growth stage (Fig. 5C), underscoring LTA’s role in directing ScaH expression and translocation.

**Figure 5.**
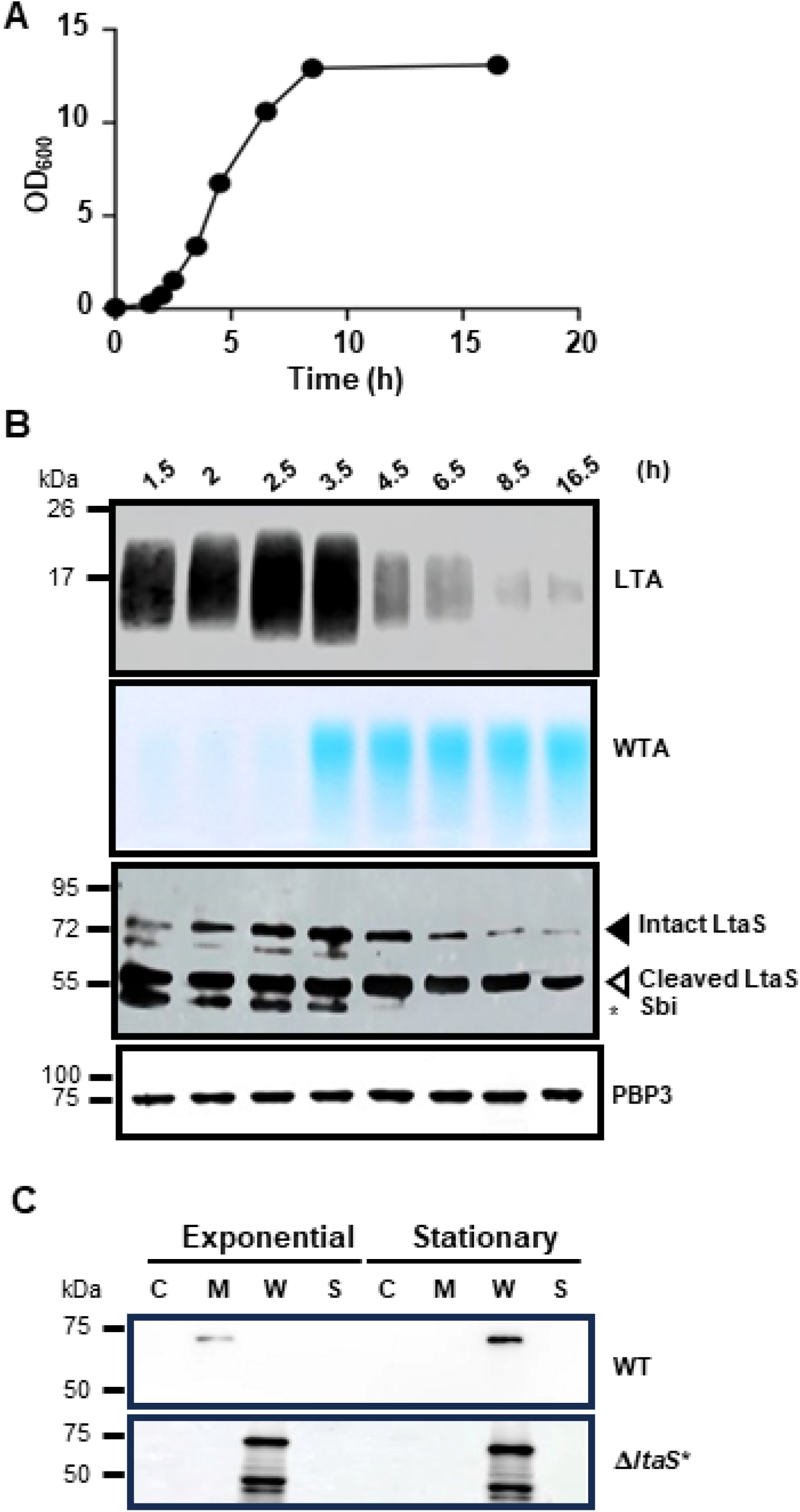
LTA determines the translocation of ScaH to the cell wall. **(A)** Growth curve of USA300 WT in TSB. Each dot represents a sampling time point. **(B)** Growth-dependent reciprocal production of LTA and WTA, and LtaS processing. LTA and LtaS were detected by Western blotting with their specific antibodies, while WTA was stained with alcian blue. PBP3 was used as a loading control. Sbi, an IgG-binding protein. **(C)** Translocation of ScaH. USA300 WT and Δ*ltaS*\* strains were grown in TSB until the exponential growth phase (t = 2 h) or stationary growth phase (t = 8.5 h). Cells were collected and fractionated, and ScaH was detected by Western blotting. C, cytoplasm; M, membrane; W, cell wall; S, culture supernatant.

### ScaH inactivation increases the cell wall integrity in *ΔltaS\**

Finally, we investigated the role of ScaH in the cell wall integrity of *S. aureus*. ScaH has the NAGase and amidase/peptidase domains and may, therefore, affect both glycan chain length and PG crosslinking. When the glycan chain length was compared between WT, Δ*ltaS*\*, *scaH*, and Δ*ltaS*:scaH* strains, the *scaH* mutation did not detectably increase the overall glycan chain length of WT or Δ*ltaS* (Fig. S10). In the PG crosslinking analysis, Δ*ltaS*\* exhibited 22-25% reduced PG crosslinking compared with WT (Fig. 6A), consistent with a previous finding (15). Although *scaH* inactivation did not detectably alter PG crosslinking in WT, it did in Δ*ltaS*\* (Fig. 6A). The Δ*ltaS*\*:*scaH* strain showed a small but significant (7% - 11%) increase in PG crosslinking compared with the Δ*ltaS*\* strain, indicating that ScaH is, at least in part, responsible for the reduced PG crosslinking in *ltaS*\*. The increased PG crosslinking in Δ*ltaS*:scaH* coincided with increased oxacillin MIC (Fig. 6B), restored osmoresistance (Fig. 6C) and resistance to oxacillin treatment in a murine model of blood infection (Fig. 6D, and S11), further demonstrating that the upregulation and uncontrolled translocation of ScaH contributes to the reduced cell wall integrity of Δ*ltaS*\*. Intriguingly, ScaH inactivation lowered the oxacillin MIC 32 – 64 times in the WT strain (WT in Fig. 6B), suggesting that its role is context-dependent.

**Figure 6.**
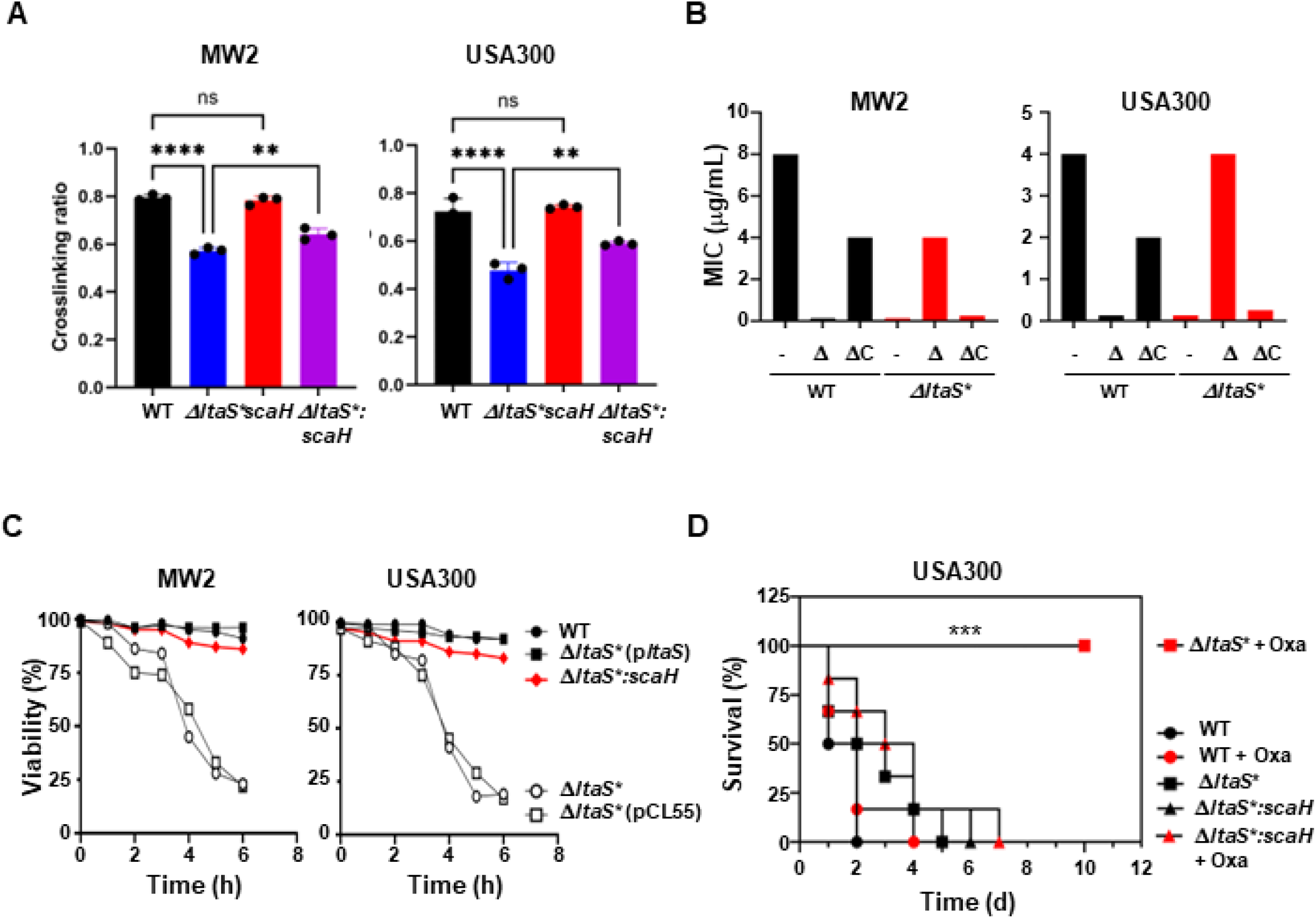
ScaH inactivation partially restores the cell wall integrity of Δ*ltaS*\*. **(A)** Effect of s*caH* inactivation on peptidoglycan crosslinking. Cell walls were purified from the test strains, and crosslinking was assessed by muropeptide analysis. The crosslinking ratio was calculated as (dimer + trimer + tetramer) / (monomer + dimer + trimer + tetramer). **(B)** Effect of *scaH* inactivation on oxacillin sensitivity of WT and Δ*ltaS*\* in TSB. MIC, minimum inhibitory concentration. -, no mutation; Δ, *scaH* inactivation; ΔC, Complemented with pCL-*scaH*. **(C)** Effect of *scaH* inactivation on osmosensitivity of WT and Δ*ltaS*\*. Cells were incubated in water for 6 h, and cell viability was analyzed with a microbial cell analyzer. p*ltaS*, pCL-*ltaS*. **(D)** Effect of *scaH* inactivation on resistance to oxacillin treatment in a murine model of blood infection. Mice were infected with the indicated strains via retro-orbital injection, followed by intramuscular oxacillin injections at 1, 4, and 8 h post-infection. Oxa, oxacillin; ***, p < 0.001 by Log-Rank test.

## Discussion

LTA was discovered over 60 years ago (26), and significant progress has been made in understanding its synthesis and physiological roles. In particular, the increased cell size, reduced PG crosslinking, increased beta-lactam sensitivity, and osmosensitivity of the Δ*ltaS* strains suggest that LTA is crucial for cell wall integrity (5, 12, 13, 15). However, the underlying molecular mechanisms have been unknown. In this study, we showed that, in *S. aureus*, LTA contributes to cell wall integrity by suppressing the expression and activity of the PGH ScaH, especially when cells are actively growing (Fig. 3 and 4). The growth-dependent expression of LTA also controls the translocation of ScaH from the membrane to the cell wall (Fig. 5).

Although not investigated further in this study, LTA binding to PBP2 and PBP4, and the partial restoration of PG crosslinking in Δ*ltaS*:scaH* imply that LTA might also contribute to the cell integrity via its interaction with those cell wall synthesis enzymes.

ScaH is a putatively bifunctional PG hydrolase predicted to have NAGase and amidase/peptidase activities (Fig. 4A) (27, 28), but its physiological functions are not known. Previous studies showed that *scaH* inactivation alone did not alter growth, cell division, lysostaphin resistance, muropeptide profile, or WTA production (28). Intriguingly, in the *ugtP* (= *ypfP*) mutant, which uses phosphatidyl glycerol, rather than diglucosyl diacylglycerol as an LTA anchor molecule and produces larger LTA, ScaH was enriched in the membrane fraction, and its secretion into the culture supernatant was reduced (29). In addition, the secretion protein profile was altered in the *ugtP* mutant (29). These findings suggest that the influence of LTA on protein translocation and secretion extends beyond the LTA-binding proteins identified in this study (Fig. 2) and that LTA may play a more extensive role in these processes than previously thought.

One of the interesting findings in this study is the reciprocal expression of LTA and WTA (Fig. 5B). LTA was abundant when cells were actively growing, whereas WTA was highly expressed during the late growth phase. Such reciprocal expression of TAs was also observed in *Streptococcus pneumoniae*. Unlike S. *aureus*, where LTA (a polymer of phosphoglycerol) and WTA (a polymer of ribitol phosphate) are structurally different and synthesized via distinct pathways, in *S. pneumoniae*, LTA and WTA have the same chemical structure and are synthesized by the same pathway, which diverges only at the last ligation step (1). The reciprocal expression of TAs in *S. pneumoniae* is due to the degradation of LtaS by FtsH at the late growth phase, switching TA synthesis from LTA to WTA (30). The TA switching also controls the membrane-to-cell wall translocation of the major PG hydrolase LytA because both TAs directly bind to LytA (30). In *S. aureus*, LtaS is cleaved and inactivated by the essential signal peptidase SpsB, not FtsH (31). Since the reduction of LTA coincided with the depletion of LtaS (Fig. 5B), it is likely that the downregulation of LTA is caused by the increased cleavage of LtaS by SpsB. Currently, it is unknown why LtaS cleavage might be enhanced during the late growth phase. It might be due to the increased expression of SpsB or the activation/stabilization of SpsB by molecules produced at the late growth phase. Since WTA production is positively controlled by the quorum sensing Agr system (32), it is possible that the WTA synthesis process might activate or stabilize SpsB through a mediator (e.g., a membrane protease), resulting in the enhanced LtaS cleavage and reduction of LTA. Nonetheless, the convergent regulatory mechanism in *S. aureus* and *S. pneumoniae* underscores how different evolutionary pressures can lead to similar functional outcomes in bacterial cell wall maintenance and adaptation.

LTA synthesis represses the transcription of *scaH* (Fig. 3). Since LTA resides outside the cell membrane, it is unlikely that LTA is directly involved in *scaH* repression. Another possibility is that LtaS, a membrane protein, might be involved in the *scaH* repression directly or indirectly through protein-protein interactions. Alternatively, the absence of LTA or the accumulation of the LTA anchor molecule Glc_2_-DAG might cause broad effects on cell physiology and envelope integrity, which can initiate complex regulatory cascades, leading to widespread transcriptional changes. Those possibilities are currently being investigated in our laboratory.

The inactivation of *scaH* caused the opposite effect on the beta-lactam resistance in WT and Δ*ltaS*.* It greatly reduced the oxacillin MIC of the WT strain but increased that of Δ*ltaS\** (Fig. 6B). It is difficult to explain why the ScaH inactivation reduces the oxacillin resistance in WT. Inactivation of ScaH could lead to structural changes in the cell wall by reducing PG turnover or altering how new cell wall material is integrated. This might cause cell wall stress that makes *S. aureus* more susceptible to beta-lactam antibiotics like oxacillin. The cell wall of *S. aureus* relies on a delicate balance of hydrolases and synthases. Inactivation of ScaH might disrupt this balance, potentially leading to weakened cell walls or structural defects. Such defects may amplify the action of oxacillin, thereby reducing the MIC.

Δ*ltaS*\* shows greatly reduced PG crosslinking compared with WT (Fig. 6A) (15). The reduced PG crosslinking can be, to a certain extent, explained by the uncontrolled release of ScaH to the cell wall during active cell growth (Fig. 3C and 5C). However, the inactivation of ScaH did not fully restore PG crosslinking in Δ*ltaS\**, suggesting an involvement of other factors. PBP2 and PBP4 might be those factors. The reduced PG crosslinking in Δ*ltaS*\* might hint at the reduced activity of PBP4, which is known to be critical for staphylococcal PG crosslinking. Alternatively, like *scaH,* the *pbp2* and *pbp4* genes might be transcriptionally regulated by LTA synthesis. Such possibilities should be further examined in future investigations.

In summary, we proposed the following model about the role of ScaH in *S. aureus* cell wall integrity (Fig. 7). In WT cells, LTA sequesters ScaH to the cell membrane during active growth and prevents premature PG hydrolysis. In the stationary growth phase, the LTA level decreases, resulting in higher expression of ScaH and translocation of the PGH to the cell wall for remodeling. In Δ*ltaS*\*, however, ScaH is constantly expressed at a higher level and translocated to the cell wall. It reduces PG crosslinking, regardless of the growth stage. As a result, the cell wall is weakened, particularly during active growth, and the cells become hypersensitive to beta-lactams and hypoosmotic conditions.

**Figure 7.**
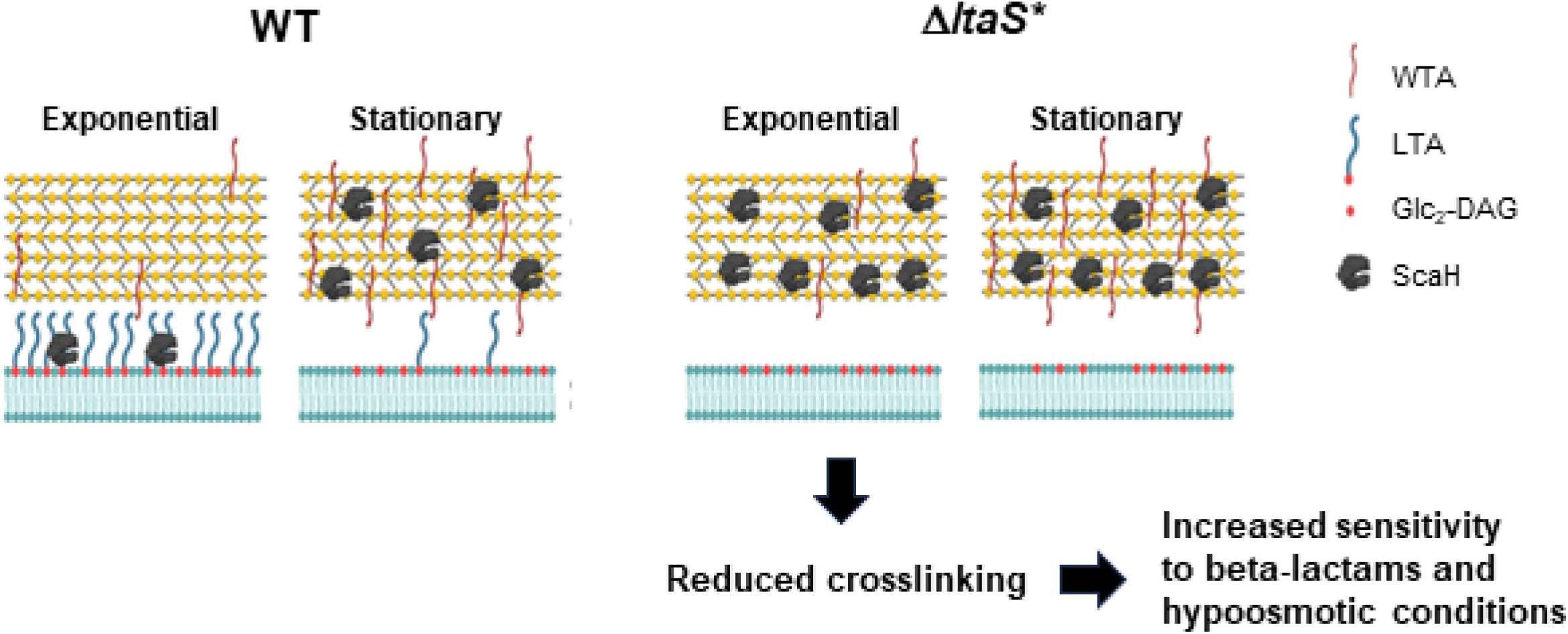
Model of LTA’s role in cell wall integrity of *S. aureus*.

## Materials and Methods

### Bacterial strains and growth conditions

The bacterial strains and plasmids used in this study are listed in Table 1. *Escherichia. coli* DH5α and BL21(DE3) were used for cloning and recombinant protein production, respectively. *E coli* was grown in lysogeny broth (LB), whereas *S. aureus* was grown in tryptic soy broth (TSB). When necessary, antibiotics were added to the growth media at the following concentrations: carbenicillin, 100 μg/mL, erythromycin, 10 μg/mL; chloramphenicol, 5 μg/mL for single-copy plasmid and 10 μg/mL for multi-copy plasmid; kanamycin, 30 μg/mL for *E. coli* and 100 μg/mL for *S. aureus*.

**Table 1.**
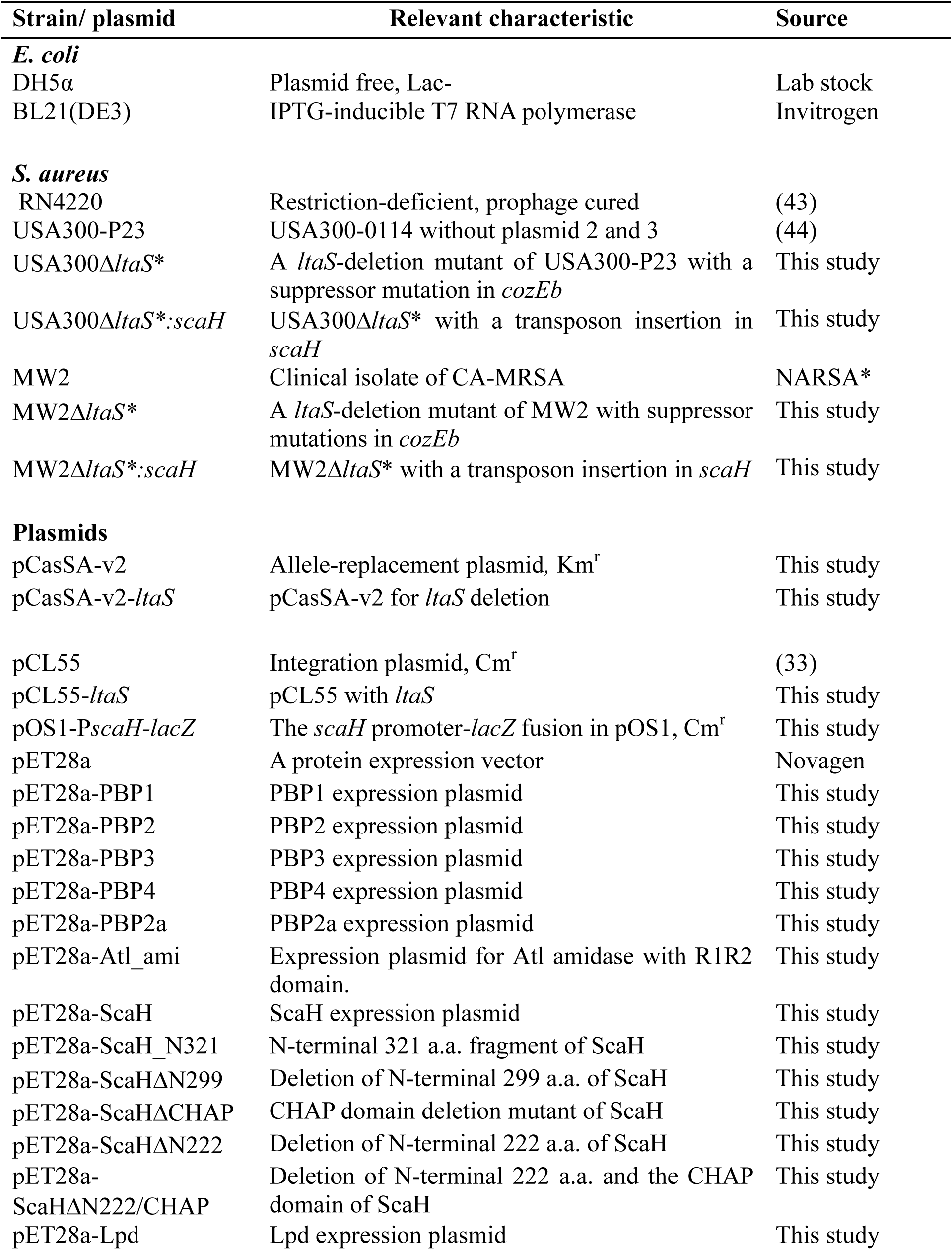
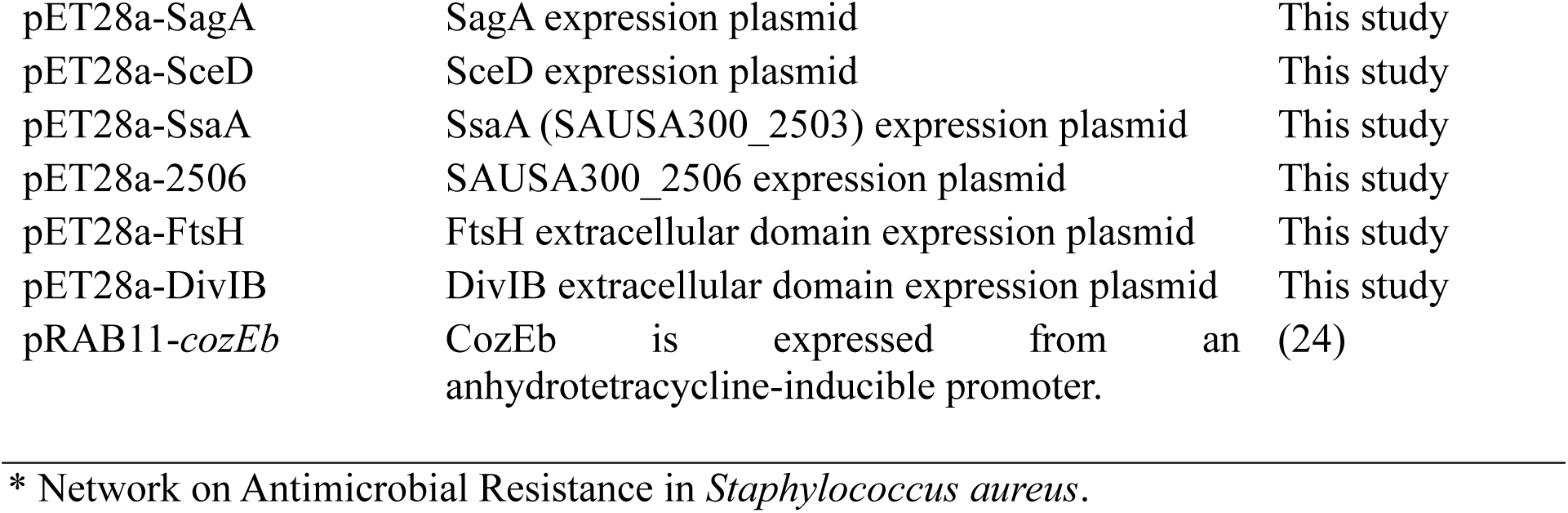
Bacterial strains and plasmids used in this study.

### Construction of plasmids

Phusion or Q5 DNA polymerase (NEB) was used for PCR amplification of DNA. To generate pCL-*ltaS*, the vector pCL55 (33) and the *ltaS* gene of USA300 were PCR-amplified with the primer sets P5661/P5662 and P5659/P5660, respectively (Table 2). All DNAs were purified and assembled with the Gibson method (34). The assembled DNA was transformed into *E. coli* DH5α. To generate pOS1-P*scaH-lacZ*, the vector was PCR-amplified with the primer set P6486/P6487 from pOS1-P*mbtS*-*lacZ* (35), while the *scaH* promoter (P*scaH*) of USA300 was amplified with the primer set P6488/P6489. The DNAs were purified, Gibson-assembled, and transformed into *E. coli* DH5a. Correct plasmids were verified by plasmid sequencing, electroporated into *S. aureus* RN4220, and then transduced into the target *S. aureus* strains with phi11 or phi85.

**Table 2.**
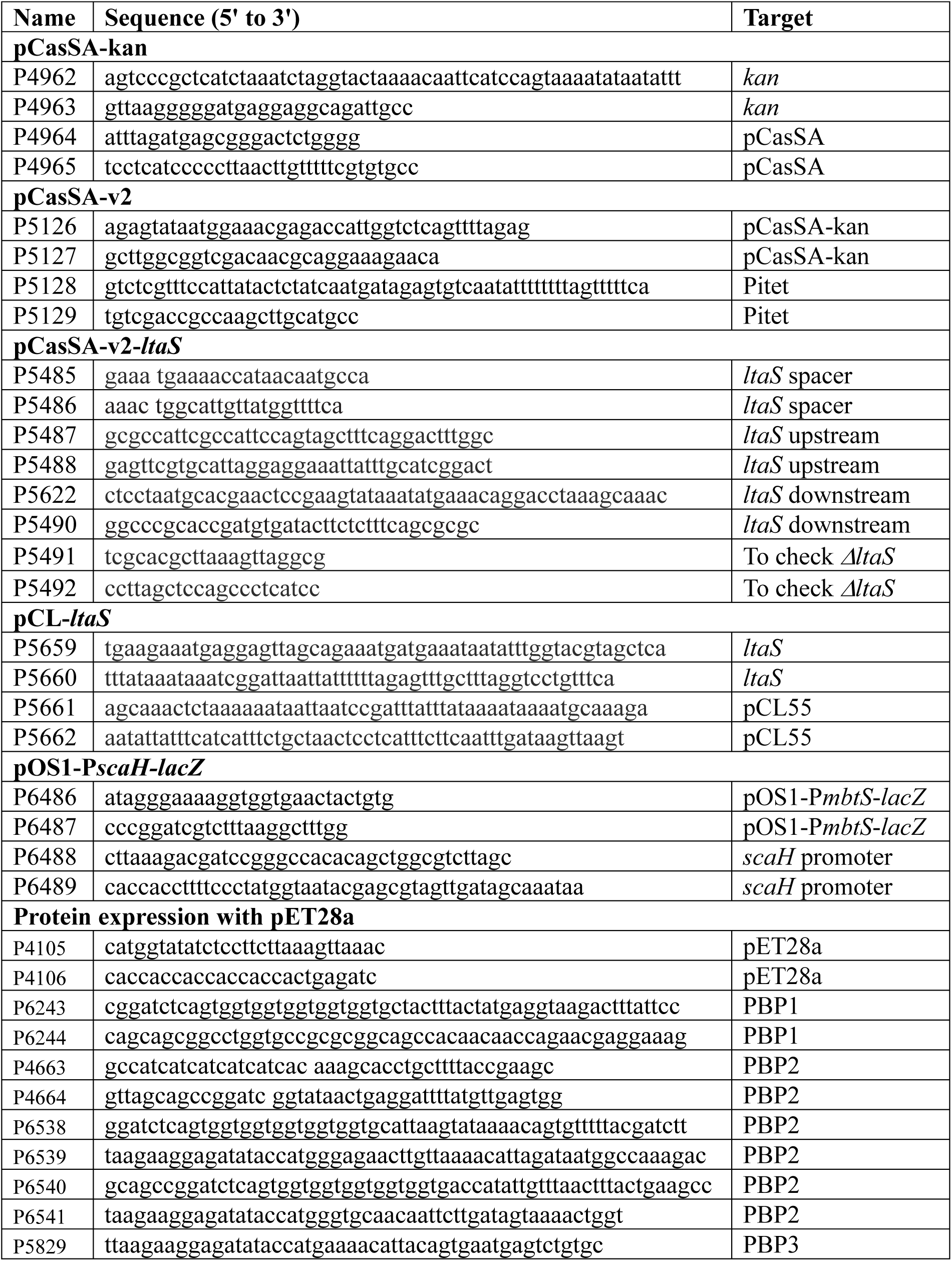
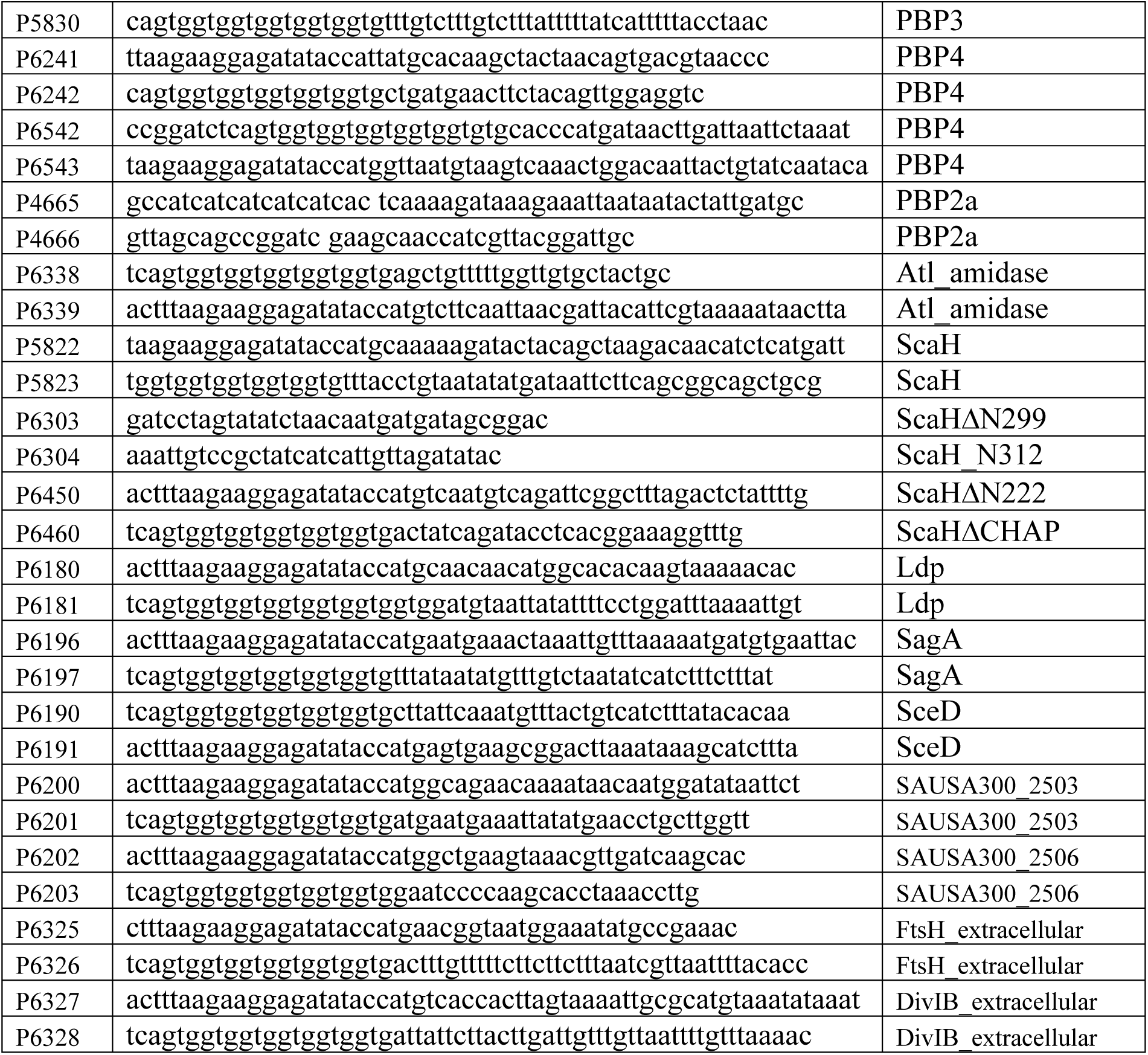
Oligonucleotides used in this study.

All recombinant proteins were expressed with C-terminal His_6_-tag with pET28a. The vector was PCR-amplified with P4105/P4106, whereas the protein sequences were PCR-amplified with the primers listed in Table 2. The PCR products were purified, and the vector and the protein genes were assembled by the Gibson method (34). The assembled products were transformed into *E. coli* DH5α. DNA sequencing confirmed the correct plasmid. Finally, the protein expression plasmids were transformed into *E. coli* BL21(DE3).

### Transmission electron Microscopy

The test strains were grown in TSB until OD_600_ of 1. Cells were collected by centrifugation and sent to the Indiana University electron microscopy facility for TEM.

### Fluorescence microscopy and cell size determination

The test strains were grown in TSB until OD_600_ of 1. Cells were collected by centrifugation, washed with PBS twice, and incubated with FITC-labeled vancomycin (1 µg/mL, Sigma) at RT for 10 min. The stained cells were imaged by a confocal microscope (FV3000, Olympus), and cell size (n = 200) was measured with Image J.

### Determination of Minimum inhibitory concentration (MIC)

MIC was determined in TSB with a microdilution method according to the recommendations of the Clinical and Laboratory Standard Institute (CLSI) (36). If necessary, MIC was also determined with MIC strips (Liofilchem). The test strains were grown in TSB to OD_600_ of 0.5, and the resulting culture (100 μL) was spread on TSA. Then, the MIC strips were placed on the plate, which was incubated at 37°C for 16 – 18 h.

### Cell viability assay

The test strains were grown to OD_600_ of 1.0. Cells were collected by centrifugation, suspended in water, and incubated for 6 h. Samples (10 μL) were taken every hour and stained with a mixture of Syto9 and PI (1:1, 10 μL, Sigma). Fig. 1G and S6 images were taken with a confocal microscope (FV3000, Olympus), whereas Fig.6C was generated with a microbial cell analyzer (Countstar Mira FL Pro, Alit Biotech).

### LTA affinity chromatography

LTA was purified from 1.5 L TSB culture of *S. aureus* USA300 WT utilizing the 1-butanol extraction method (37). The purified LTA was coupled to epoxide resin (Profinity Epoxide Media, Bio-Rad), and the free active groups on the resin were blocked with 1 M ethanolamine (pH 8). The coupled resin was sequentially washed with an acidic buffer (100 mM acetate, 500 mM NaCl, pH 4.0) followed by a basic buffer (100 mM phosphate, 500 mM NaCl, pH 8.0). An alanine column was generated with alanine in the same manner for a control.

To identify LTA-binding proteins, membranes were fractionated from *S. aureus* USA300 WT (35) and suspended in the binding buffer (20 mM Tris HCl pH 8, 50 mM NaCl, 0.05% Triton X-100). The membrane suspension was passed through the LTA and the alanine columns, and the columns were washed with the binding buffer. The binding proteins were eluted with the elution buffer (20 mM Tris-Cl pH 8, 500 mM NaCl, 0.05% Triton X-100). The eluted fractions were analyzed by SDS-PAGE and silver staining. Protein bands present only in the LTA column were cut out from the gel and sent to the Laboratory for Biological Mass Spectroscopy at Indiana University for mass spectrometry.

### Protein expression and purification

*E. coli* BL21(DE3) carrying the target plasmid was grown in LB at 30°C to OD_600_ ∼0.6, and the protein expression was induced with 1 mM IPTG for 6 h. Bacterial cells were lysed with lysozyme treatment and sonication. Cell lysates were centrifuged (10 k ×g, 30 min), and the supernatant was subjected to Ni-NTA column chromatography. The proteins bound to the column were eluted with 0.15 M imidazole. The eluted proteins were dialyzed into 50 mM Tris HCl, pH 7.6, 0.1M NaCl, quantified by Nanodrop at 280 nm, and stored frozen at −80°C until use.

### ELISA

We carried out the ELISA assay as described previously (38). Briefly, a 96-well plate (Corning) was coated with test proteins (100 µL, 1 mg/mL) in PBS overnight at 4°C, washed, and blocked. Then, varying amounts of LTA (0 – 300 ng/µL) in PBS were added and incubated at 37°C for 4 h. Bound LTA was measured with anti-LTA antibody (Clone 55, HyCult Biotechnology). and HPR-conjugated anti-mouse antibody (Santa Cruz).

### Western blotting

Western blot analysis was performed as described previously (39). All primary antibodies were generated by our laboratory except for the LTA antibody. The Western blots were visualized by the Supersignal West Pico PLUS Chemiluminescent Substrate (Thermo Scientific), and the images were taken and processed with AZI400 (Azure Biosystems).

### WTA detection

WTA extraction and detection were carried out by following the methods described by Karinou et al (15). Cells were normalized by OD_600_.

### RBB labeled peptidoglycan degradation assay

Purified peptidoglycan (PG) was labeled with Remazol Brilliant Blue (RBB) as described (25). Labeled PG (10 μL) was mixed with 90 μL PBS and incubated at 37°C for 30 min. ScaH (10 nM), with or without LTA (100 μg), was added, and the samples were further incubated at 37°C. At 2, 4, and 8 h, ethanol was added, and samples were centrifuged at 400,000 ×g for 30 min at 25°C. The supernatants were collected, and OD_595_ was measured.

### ScaH localization test

Overnight cultures of WT and Δ*ltaS*\* strains of MW2 and USA300 were diluted 100 times in 3 mL TSB and incubated at 37°C, 200 rpm for 2 h (early exponential growth phase) or for 8.5 h (early stationary growth phase). Cells were collected by centrifugation and fractionated as described previously (35). The fractionated samples were subjected to 10% SDS-PAGE and Western blotting with anti-ScaH mouse antibodies at 1:10,000 dilution and HRP-linked anti-mouse IgG antibody (Santa Cruz) at 1:20,000 dilution.

### Muropeptide analysis

Muropeptides were extracted from *S. aureus* following a protocol with some modifications (40, 41). Briefly, overnight cultures were grown to OD_600_ 0.7, and sacculi were prepared. After WTA removal, sacculi were digested with mutanolysin (4,000 U/mL) in NaH_2_PO_4_ buffer (pH 5.5) at 37°C for 16 h. Lyophilized digests were resuspended in water, and 15 μL was injected for LC-MS analysis.

LC-MS separation and analysis followed established methods (42). Extracted ion counts (EIC) for monomer (1253.5856 [M+H]^+^), dimer (1209.0617 [M+2H]^2+^), trimer (1194.2204 [M+3H]^3+^), and tetramer (1186.7998 [M+4H]^4+^) were obtained. The crosslinking ratio = (dimer + trimer + tetramer) / (monomer + dimer + trimer + tetramer).

### Animal test

The test strains were grown in TSB at 37°C until OD_600_ of 1. Cells were collected, washed with PBS, and suspended in PBS to OD_600_ 8. The bacterial cells (50 μL) were administered into six C57BL/6J mice (3M and 3F) via retro-orbital injection. At 1, 4, and 8 h post-infection, the mice were i.m. injected with oxacillin (100 μL, 20 mg/mL) and observed for 2 weeks.

The animal experiment followed the Guide for the Care and Use of Laboratory Animals of the National Institutes of Health. The animal protocol NW-48 was approved by the Committee on the Ethics of Animal Experiments of the Indiana University School of Medicine-Northwest. Every effort was made to minimize the suffering of the animals.

### Statistical analysis

Statistical significance was assessed with Prism 10 (GraphPad).

## Acknowledgments

We thank Morten Kjos at the Norwegian University of Life Sciences for providing pRAB11-*cozEb* and for his valuable feedback on the manuscript. This study was supported by NIH (AI143792) to TB and AI148752 to SW. The funders had no role in study design, data collection, interpretation, or the decision to submit the work for publication.

## Author Contributions

AK, MS, YP, and TB conceived and designed the experiments. AK, MS, YP, BJ, PAL, MB, EMF, AS, and DS performed the experiments. AK, MS, YP, PL, SW, and TB analyzed and interpreted the data. AK, MS, YP, and TB prepared figures and tables. AM, YP, and TB wrote the first draft of the manuscript. All authors reviewed and contributed to the manuscript and approved its final version.

